# Towards CRISPR/Cas9-based gene drive in the diamondback moth *Plutella xylostella*

**DOI:** 10.1101/2021.10.05.462963

**Authors:** X. Xu, T. Harvey-Samuel, H. Siddiqui, J. Ang, M.A.E Anderson, C. Reitmayer, E. Lovett, P.T. Leftwich, M. You, L. Alphey

## Abstract

Promising to provide powerful genetic control tools, gene drives have been constructed in multiple dipterans, yeast and mice, for the purposes of population elimination or modification. However, it remains unclear whether these techniques can be applied to lepidopterans. Here, we used endogenous regulatory elements to drive Cas9 and sgRNA expression in the diamondback moth, (*Plutella xylostella*), and test the first split-drive system in a lepidopteran. The diamondback moth is an economically important global agriculture pest of cruciferous crops and has developed severe resistance to various insecticides, making it a prime candidate for such novel control strategy development. A very high level of somatic editing was observed in Cas9/sgRNA transheterozygotes, although no significant homing was revealed in the subsequent generation. Although heritable, Cas9-medated germline cleavage, as well as maternal and paternal Cas9 deposition was observed, rates were far lower than for somatic cleavage events, indicating robust somatic but limited germline activity of Cas9/sgRNA under the control of selected regulatory elements. Our results provide valuable experience, paving the way for future construction of gene drive-based genetic control strategies in DBM or other lepidopterans.

## Introduction

Gene drives are heritable elements capable of autonomously increasing their frequency within a gene pool^1,2^. Traits associated with the gene drive will also spread and could be arranged to include examples beneficial for pest control (e.g. a sex-specific fitness cost), reducing vector-borne virus transmission (e.g. virus-refractory transgenes) or conservation (e.g. resistance to disease/pesticides)^3,4^.

One example of a gene drive technology is the CRISPR-Cas9 based homing drive^4,5^. In its simplest form, this system requires a source of Cas9 and a sgRNA expression cassette to be integrated into a genome at the precise site specified by the sgRNA. As this integrated ‘homing element’ has disrupted its linked sgRNA site, it is immune to further cutting. However, in a heterozygote, the homologous ‘wild-type’ chromosome may be targeted for cleavage by Cas9 and in the resulting DSB repair, the locus harbouring the integrated homing element may be used as a template for homology-directed repair (HDR), transforming the heterozygous cell into one homozygous for the homing element (a process known as ‘homing’). If this process occurs in the germline, it can bias the inheritance of the homing element above the expected 50:50 ratio^5^.

The binary nature of the CRISPR/Cas9 system allows variations on this form, e.g. with one or both components integrated at sites away from the homing element. Such ‘split-drive’ (sgRNA cassette remains in homing element)^6–9^ or ‘trans-drive’ (neither component remains in homing element)^10^ designs are predicted to act in a more ‘controllable’ way once released into a target population as any component not included in the homing element will not benefit from super-Mendelian inheritance and will likely reduce in frequency over time due to associated fitness costs^11,12^. As both components are required in an individual for the drive to function, the efficiency of the drive will thus reduce over time as one or more components become limiting. Although potentially less efficient than the simple ‘all in one’ design outlined above, the potential to geographically and temporally limit the spread of a gene drive may prove beneficial to the regulation, perception and field deployment of such technologies^13^.

To date, CRISPR/Cas9 homing drives have been demonstrated in various dipterans (prominent examples including *Drosophila melanogaster*^14^, the mosquitoes *Anopheles gambiae*^15^, *Anopheles stephensi*^16^ and *Aedes aegypti*^17^), yeast^6^ and mice^18^. Here we explore the potential of such technology for the first time in a lepidopteran – the diamondback moth (DBM) *Plutella xylostella* – focussing on a ‘split-drive’ design. Lack of previous research into lepidopteran gene drive is surprising, given the extreme and wide-ranging importance of this insect order, e.g. as major agricultural pests and invasive species^19^, and as producers of silk and components of industrial processes^20^. DBM provides an appropriate model for such work as it is both a globally important pest of brassica crops^21,22^, and is tractable in terms of genetic engineering technologies, having been the subject of previous genetics-based pest control strategy development^23,24^.

## Materials and methods

### Insects

DBM transgenic lines were generated from the Vero Beach wildtype strain. Rearing conditions were as described previously^25,26^.

### Identification and expression profiling of germline candidate genes

To drive Cas9 expression in specific tissues and developmental stages, germline-active promoters must be identified and characterized. Referring to previous reports ^27–30^, amino acid sequences of eight germline candidates *BGCN*, *Shutdown*, *SDS3*, *Meiw68*, *SIWI*, *NanosO*, *NanosP* and *NanosM* (origins listed in Table S2) were downloaded from NCBI (https://www.ncbi.nlm.nih.gov/) and blasted against DBM pacbiov1 genome database (http://ensembl.lepbase.org/Plutella_xylostella_pacbiov1/Info/Index) to identify putative homologs.

Expression patterns of these DBM candidates were compared with RT-PCR. DBM samples collected at different developmental stages (including embryonic, larval, and adult stages) were dissected, frozen in liquid nitrogen, placed in RNAlater (Thermo Fisher Scientific) and stored at −80 °C. Total RNA was extracted with RNeasy Mini Kit (Qiagen), followed by removal of gDNA with DNase I (Thermo Fisher Scientific). Purified RNA was diluted to 20 ng/μl and used to produce cDNA pools with RevertAid First Strand cDNA Synthesis Kit (Thermo Fisher Scientific). RT-PCR was conducted using the Q5 High-Fidelity 2X Master Mix (NEB). All primers used in the current study are available in Table S1.

### Construct design

Adult ovary total RNA was used to amplify the 5’ and 3’ UTR of *Pxmeiw68* and *PxnanosP* using the SMARTer RACE 5’/3’ Kit (Takara, Japan). 5’ and 3’ regulatory sequences were then cloned from 4^th^ instar larval gDNA, (NucleoSpin Tissue Column - Macherey-Nagel). *PiggyBac* constructs AGG1906 (*Pxmeiw68-Cas9* Genbank accession: OK145566) and AGG2093 (*PxnanosP-Cas9* Genbank accession: OK145567) were developed as for AGG1536 (*Pxvasa-Cas9*) previously^25^, except for replacing the *Pxvasa* promoter, *Pxvasa* 3’UTR and SV40 terminator with corresponding regulatory fragments from *Pxmeiw68* and *PxnanosP* as well as the fluorophore marker 3’ UTR with P10. All plasmids were assembled with NEBuilder HiFi DNA Assembly Cloning Kit (NEB). See Figure 1A.

**Figure 1.**
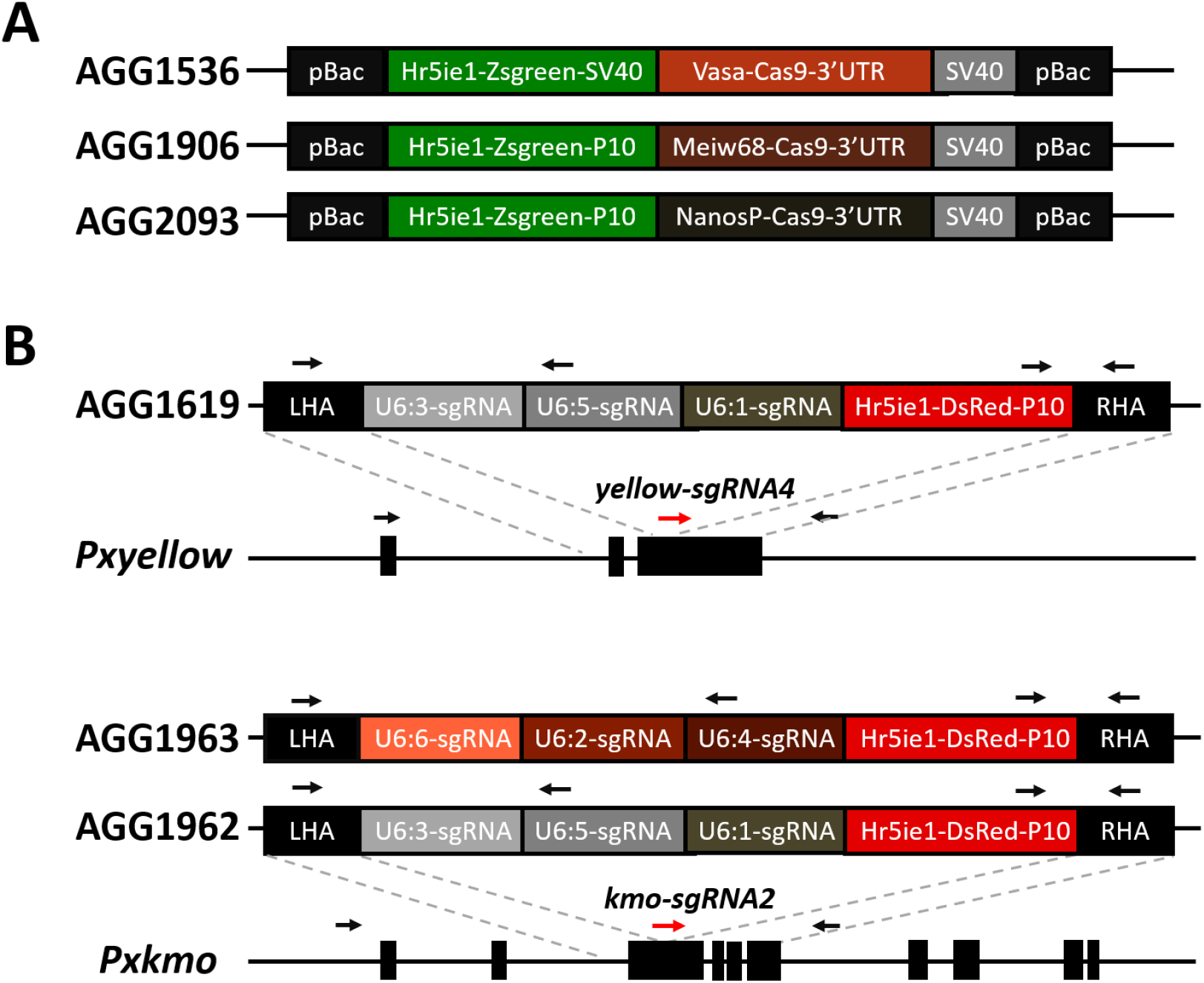
Construction of transgenic lines. A: Donor cassettes for *piggyBac*-mediated transformation of Cas9 lines. B: HDR-based integration of homing elements into endogenous marker genes *Pxyellow* and *Pxkmo*.

For constructing sgRNA expressing lines (‘homing elements’), two pigmentation genes *Pxyellow* and *Pxkmo*^26^ were selected as targets for CRISPR-based site-specific insertion (‘knock-in’) of sgRNA cassettes, due to the ease of visually scoring null mutations in these genes. 800~1000bp upstream/downstream regions immediately flanking each sgRNA cleavage site were cloned from gDNA as homology arms for mediating HDR repair. Six RNA polymerase III *PxU6* promoters^31^, were divided into two sets of three and used to express relevant sgRNAs for each gene (see Figure 1B). sgRNAs were designed with CHOPCHOP (https://chopchop.cbu.uib.no/). sgRNA expression cassettes were synthesised and cloned into the homology arm destination plasmid, alongside a Zsgreen-expressing marker (Figure 1B) (Genewiz). (Genbank accession numbers AGG1619:OK145568, AGG1962:OK145570, AGG1963:OK145569).

Assembled constructs were purified with NucleoBond Xtra Midi Prep Kit EF (Macherey-Nagel) before injection.

### Development of transgenic lines

A *Pxvasa-Cas9* line (1536A) was built previously^25^. To mitigate issues associated with “positional effects” of the transgene insertion site, additional *Pxvasa-Cas9* lines were generated here, initially using the same method as previously described. Subsequently, to improve transformation efficiency, *piggyBac* helper plasmid and codon-optimized mRNA were replaced with a commercially synthesised *piggyBac* mRNA (TriLink Biotechnologies, UK). Constructs (AGG1906 or AGG2093) were injected at 500 ng/μl alongside 600 ng/μl *piggyBac* mRNA to generate *Pxvasa-Cas9*, *Pxmeiw68-Cas9* and *PxnanosP-Cas9* lines.

For constructing homing elements, sgRNAs were *in vitro* transcribed (150 ng/ul), complexed with 300 ng/ul Cas9 protein^26^ and injected alongside the relevant sgRNA plasmid (800 ng/ul). Injection, transformant and insertion-site identification were as previous^26,32^. For resolving the 5’ junction between the integrated homology arm and the genomic flanking sequence in 1619P15, the Cas9 Sequencing Kit (Oxford Nanopore Technologies, UK) was used, as PCR could not confirm the junction. In brief, high molecular weight genomic (HMW) gDNA was extracted from homozygous 1619P15 pupae using the Monarch HMW DNA extraction kit for tissue (NEB). The Cas9 Sequencing Kit (Oxford Nanopore Technology) was used to prepare a library targeting the insertion. Targets GAACTCGGTGATGACGTTCT<u>CGG</u> and GCTGAAGGGCGAGACCCACA<u>AGG</u> were designed to target the DsRed ORF using CHOPCHOP and synthesized as Alt-R CRISPR-Cas9 crRNAs (IDT). These along with the Alt-R CRISPR-Cas9 tracrRNA and Alt-R SpCas9 Nuclease V3 (IDT) were used to digest the HMW gDNA following the manufacturer’s instructions. Library was run on a MinION for 72 hours with default parameters. Epi2me was used to map the resulting FASTQ files to the AGG1619 plasmid sequence as a reference, with 308 reads aligning. Further analysis was performed using Galaxy^33^. Porechop (Version 0.2.4, https://github.com/rrwick/Porechop) was used to trim adapters from all FASTQ files, all files were compiled into a single FASTA and filtered based upon alignment to the 5’ end of the homology arm in the reference sequence using Biopython^34^. The 46 remaining reads were manually mapped to the construct/genome interface.

The *Pxkmo* knockout line was generated previously^26^. The *Pxyellow* knockout line was generated following previous protocols^35^, except using a newly designed sgRNA (*Pxyellow-sgRNA4*) targeting the third *Pxyellow* exon.

### Assessment of Cas9 somatic editing and homing efficiency

To assess somatic editing efficiency, reciprocal crosses were conducted using Cas9 and sgRNA lines. In each cross replicate, four Cas9 heterozygote ‘grandparents’ and four sgRNA heterozygote ‘grandparents’ were crossed, with F_1_ eggs collected and reared until pupation. F_1_ pupae were screened for fluorescence and body/eye pigmentation patterns, based on mutant phenotypes reported previously^26,35^. Note that *Pxyellow* mosaics are only observable when they co-occur in an individual displaying the ‘stripy’ phenotype. This phenotype occurs naturally and consists of two darkly pigmented areas running laterally along the dorsal surface of the pupae. Therefore, the number of *Pxyellow* mosaics was calculated against the number displaying this phenotype, rather than the total number of pupae. The mutagenesis of target sites was confirmed in a subset of somatic mosaics using a T7 endonuclease assay (NEB, UK).

To assess ‘homing’ activity, transheterozygous F_1_s containing both Cas9 and sgRNA transgenes (parents) were selected from the above cross and themselves crossed to *Pxyellow* or *Pxkmo* knockout lines, depending on which sgRNA line was used in the initial cross (Figure 3). In each cross replicate, a minimum of three transheterozygotes and three knockout individuals were reciprocally crossed, with five replicates conducted for each combination. An exception to the above process was made for the 1906 lines where, due to the large number of lines to assess, the ‘grandparent’ factor was not applied. In all cases, the 3^rd^ instar larvae (*Pxyellow* cross) or pupae (*Pxkmo* cross) of F_2_ progeny were screened for fluorescence (to assess gene drive activity) and knockout phenotypes (to assess germline/somatic editing efficiency). Control crosses were conducted by reciprocally crossing each sgRNA line as heterozygotes to its respective knockout line and scoring the same details as above.

### Statistical analyses

Analyses of sgRNA cassette inheritance was conducted using R version 3.6.2^36^ and figures generated using ‘ggplot2’^37^ and ‘patchwork’^38^. The proportion of offspring inheriting the sgRNA cassette was analysed using a binomial GLM (‘logit’ link) with control cross outcomes set as the intercept. Approximate 95% confidence intervals were calculated using the Sidak adjustment for multiple comparisons.

Figures of germline cleavage percentages were made using the Graphpad software. Differences were analysed with one-way ANOVA or independent *t*-test in using SPSS.

### Western blot

Moth carcass, ovary and testis samples from 2-3 day old adults were homogenised on ice in 100 μl 1x passive lysis buffer (PLB) including Halt Protease Inhibitor Cocktail (Thermo Scientific). After 15min of incubation at RT, samples were centrifuged for 10 min at 10,000 g and 4°C. An additional 50 μl of PLB + protease inhibitor was added to the samples and incubated for further 15 min at room temperature.

Protein concentrations were determined using the DC Protein Assay (BioRad) using a Bovine Serum Albumin standard curve. Absorbance values were obtained using a Microplate reader (GloMax) at 750 nm.

For each sample, equal amounts of total protein (35μg) were combined with Laemmli sample buffer (BioRad) and denatured for 5 min at 95°C.

Samples were loaded onto 4–20% precast polyacrylamide gels (BioRad) and run at 200V. A Chemiluminescent Protein Ladder (8-260kDa, LI-COR) was loaded alongside the protein samples to estimate the position of the 158kDa Cas9 (FLAG-tagged) band. Proteins were transferred onto a nitrocellulose membrane (BioRad). After transfer, unspecific binding sites were blocked over night at 4°C with 3% milk powder (Marvel) in Tris Buffered Saline solution (BioRad) including 1% Tween 20 (Sigma, TBS-Tween). Anti-FLAG-tag antibody (Abcam) was diluted 1:1000 in 3% milk/TBS-Tween and the membrane incubated over night at 4°C. Membranes were developed using Pierce ECL Western Blotting Substrate (Thermo Scientific) and scanned using a Gel Doc imager (BioRad) and analysed using Image Lab software (BioRad).

## Results

### Cas9 line generation and characterisation

Previously, we showed that *Pxvasa* was preferentially expressed in adult soma and gonads, and the selected *Pxvasa* promoter fragment drove Cas9 somatic and gonadal transcription in a single generated line (1536A)^25^. Here, three additional 1536 lines were developed (Figure 1, Table 1). Moreover, homologs of eight putative germline expressing genes were identified in DBM (Table S2), with RT-PCR showing that detectable *Pxmeiw68* expression was restricted to adult gonads while *PxnanosP* was preferentially detected in adults (especially female ovaries) (Figure S1A). The putative promoters and 5’ and 3’ regulatory regions of these two genes were cloned to direct Cas9 expression with seven 1906 (*Pxmeiw68-Cas9*) and five 2093 (*PxnanosP-Cas9*) lines subsequently generated (Table 1). Flanking PCR confirmed independent insertion sites of each line (Table S3). Paired expression profiling of Cas9 and endogenous genes (*Pxmeiw68*, *Pxvasa* and *PxnanosP*) showed that Cas9 mRNA was transcribed in both testes and ovaries of all 1536, 2093 and 1906 lines except 1906B (Figure S1B). Unexpectedly, it was also observed that expression levels of Cas9 were much higher in adult carcasses than in gonads - the opposite trend for the respective endogenous genes (Figure S1B).

**Table 1.**
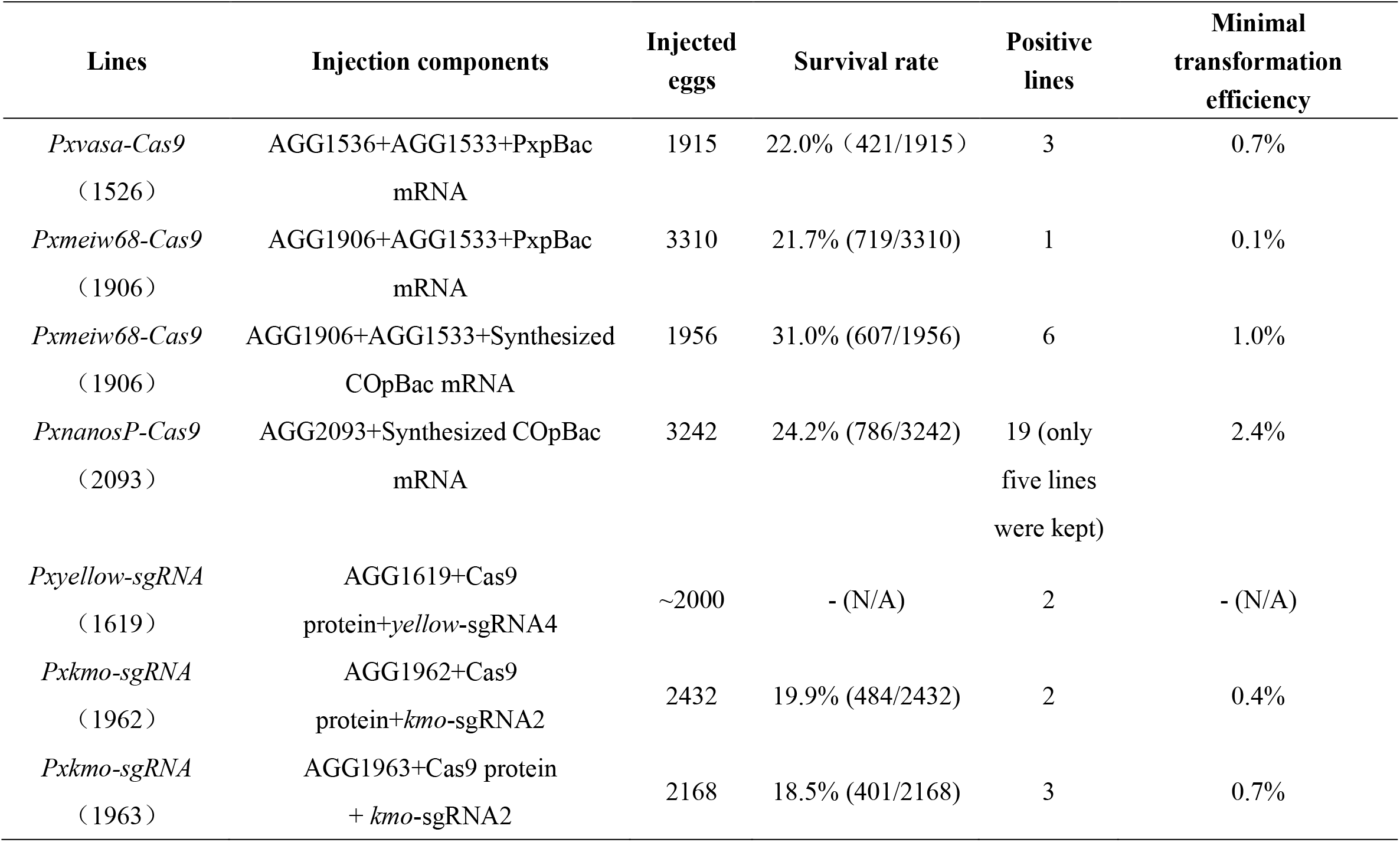
*PiggyBac*-based construction of the *Pxmeiw68-Cas9* and *PxnanosP-Cas9* transgenic lines.

### sgRNA line generation and characterisation

Homozygous null-mutations in *Pxyellow* or *Pxkmo* cause visible phenotypes in the body or eye pigmentation of DBM, respectively, making them scorable markers for genome editing events^26,35^. As such we chose these two genes as targets for building sgRNA expression lines. Although six *PxU6* promoters (*PxU6:1* to *PxU6:6*) have been identified and characterized in DBM somatic cells^31^, no information is available on their germ-cell activities. Therefore, we employed a conservative approach, utilising all six in two separate arrays (*PxU6:3,5,1* and *PxU6:6,2,4*) to express sgRNAs in three knock-in constructs (Figure 1B). In each construct the three chosen promoters each expressed an identical sgRNA, which was the sgRNA used to target the construct to its genomic location. Using CRISPR-mediated site-specific knock-in, two AGG1619 lines (*yellow-sgRNA4* expressed by *PxU6:3,5,1* promoters), two AGG1962 lines (*kmo-sgRNA2* expressed by *PxU6:3,5,1* promoters) and three AGG1963 (*kmo-sgRNA2* expressed by *PxU6:6,2,4* promoters) lines were isolated (Table 1). Interestingly, PCR analysis of the genomic location of these lines revealed a high number of non-canonical repair events (Figure S2). Of the seven insertions, two were deemed ‘off-target’ judging by the inability to generate PCR flanking bands and the lack of expected knockout phenotype in these lines when inbred (1962B, 1963A): two were in the correct locus but showed truncations of the outer transgene/homology arm sequences or genomic flanking regions (1619T18, 1962A) and three were perfectly integrated but displayed no (1619P15) very mild (1963C) or severe (1963B) internal deletions within the transgene sequence. Note that the 5’ junction of 1619P15 could not be resolved by PCR and required targeted sequencing of the region to identify the repair junction (Cas9 Sequencing Kit (Oxford Nanopore Technology) (Figure S2/S3). As 1619P15, 1962A and 1963C were the most intact, each possessing a different sgRNA expression cassette, and at least one perfectly repaired flanking sequence (both junctions perfectly repaired in 1963C and 1619P15), they were maintained for further cross experiments.

### Somatic activity of CRISPR/Cas9 system (F_1_ analysis)

To test the CRISPR/Cas9 system’s ability to function when engineered into DBM, we first crossed heterozygous Cas9 and sgRNA F0s and analysed the phenotypes and genotypes of their F_1_ progeny with regards to the target locus (*Pxyellow* or *Pxkmo*, respectively). Regarding insertion sites, these crosses theoretically resulted in four equally represented genotypes - *Cas9*-only, *target^sgRNA^* only, *Cas9* + *target^sgRNA^* transheterozygotes and non-fluorescent wildtype (WT).

#### Pxyellow analysis

Broadly, *Pxyellow* somatic mosaics were observed in *Cas9* + *yellow^sgRNA^* transheterozygotes in all crosses involving 1536 and 2093 lines, while only two out of seven 1906 lines (1906B and 1906E) showed observable mosaicism in F1s (Figure 2A). Choosing a subset of these lines for finer-scale analysis, we calculated the somatic editing rates of F_1_ individuals using 1536C, 1906B, 1906E and 2093A (Table 2). 100% of 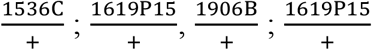 and 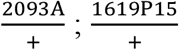 were mosaic, while lower levels (72.7-85.7%) observed in 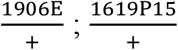 transheterozygous pupae. No mosaicism was observed in F_1_s other than transheterozygotes, with the exception of *yellow^sgRNA^*-only individuals which had inherited Cas9 from their female 2093A parent (27.2% mosaics), implying deposition of Cas9 through the maternal germline into the developing embryo in that line.

**Figure 2.**
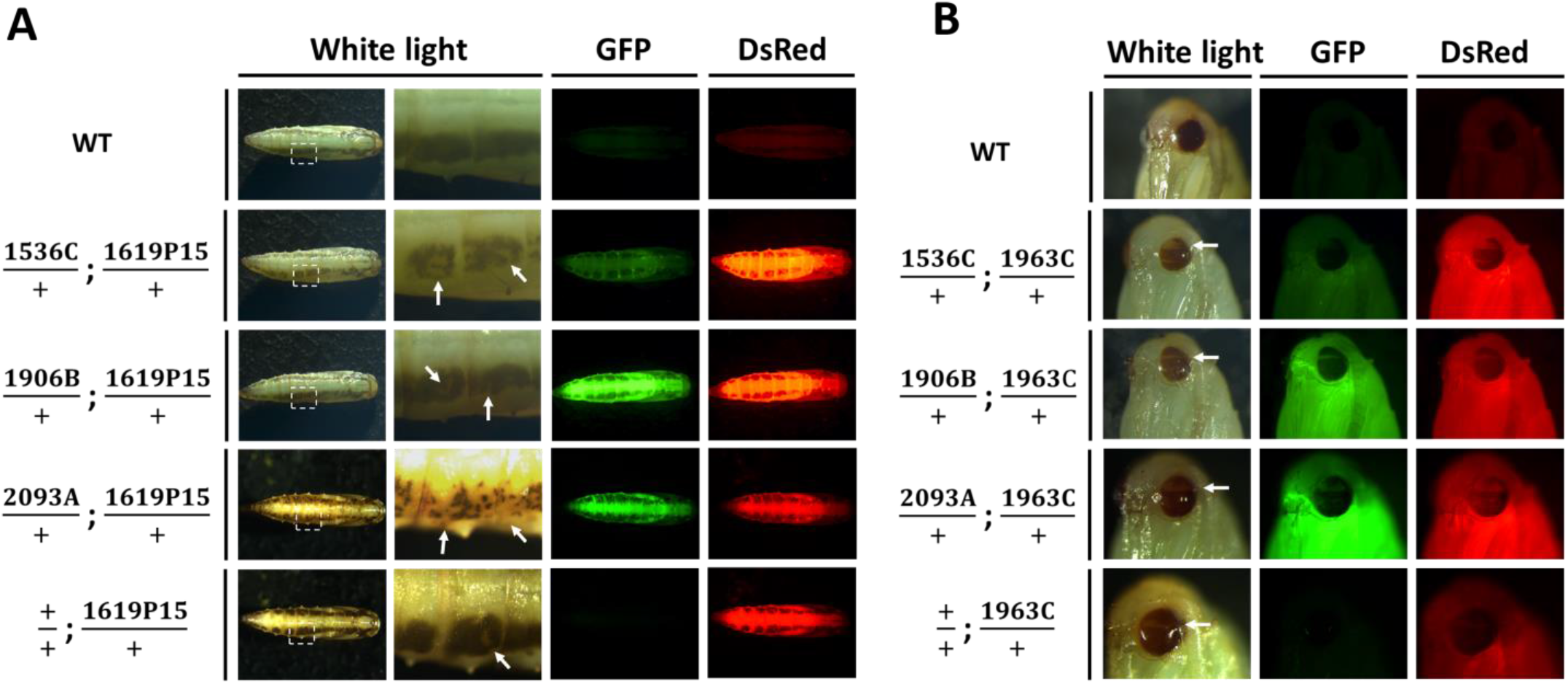
Somatic mosaicism present in F_1_ transheterozygotes. A: Parental Cas9 lines were crossed with the 1619P15 line integrated into and expressing sgRNAs targeting *Pxyellow*. B: Parental Cas9 lines are crossed with the 1963C line integrated into and expressing sgRNAs targeting *Pxkmo*. Mosaic phenotypes are indicated using white arrows.

**Table 2.** Individuals and percentages of different genotypes and phenotypes shown in F_1_s generated from *Pxyellow* homing crosses.

| Genotype | *yellow<sup>sgRNA</sup>* only |  |  | *Cas9* + *yellow<sup>sgRNA</sup>* |  |  | *Cas9* only |  |  | Non-fluorescence |  |  |
| --- | --- | --- | --- | --- | --- | --- | --- | --- | --- | --- | --- | --- |
| Phenotype | Total<br>pupae | Stripy<br>pupae | Mosaic<br>stripy<br>pupae | Total<br>pupae | Stripy<br>pupae | Mosaic<br>stripy<br>pupae | Total<br>pupae | Stripy<br>pupae | Mosaic<br>stripy<br>pupae | Total<br>pupae | Stripy<br>pupae | Mosaic<br>stripy<br>pupae |
| 1906B ♂ x 1619P15 ♀ | 60 | 24 | 0 | 58 | 16 | 16 (100.0%) | 80 | 30 | 0 | 73 | 27 | 0 |
| 1906B ♀ x 1619P15 ♂ | 62 | 30 | 0 | 45 | 17 | 17 (100.0%) | 61 | 30 | 0 | 68 | 27 | 0 |
| 1906E ♂ x 1619P15 ♀ | 99 | 13 | 0 | 105 | 14 | 12 (85.7%) | 77 | 7 | 0 | 100 | 27 | 0 |
| 1906E ♀ x 1619P15 ♂ | 51 | 17 | 0 | 40 | 11 | 8 (72.7%) | 28 | 7 | 0 | 45 | 14 | 0 |
| 1536C ♂ x 1619P15 ♀ | 56 | 8 | 0 | 63 | 7 | 7 (100.0%) | 58 | 18 | 0 | 52 | 16 | 0 |
| 1536C ♀ x 1619P15 ♂ | 35 | 12 | 0 | 64 | 16 | 16 (100.0%) | 46 | 19 | 0 | 57 | 16 | 0 |
| 2093A ♂ x 1619P15 ♀ | 97 | 59 | 0 | 83 | 47 | 47 (100.0%) | 72 | 40 | 0 | 86 | 45 | 0 |
| 2093A ♀ x 1619P15 ♂ | 98 | 81 | 22 (27.2%) | 92 | 84 | 84 (100.0%) | 119 | 98 | 0 | 107 | 82 | 0 |

#### Pxkmo analysis

Regarding crosses involving the 1963C line, *Pxkmo* mosaicism broadly concurred with *Pxyellow* results above. At finer scale, however, the observable mosaic phenotype was generally lower in 1536C (54.4-84.3%), 1906B (4.0-9.7%) and 1906E (0-1.8%) F_1_ transheterozygotes, although 100% of 2093A transheterozygotes continued to show some form of eye mosaicism (Table 3). Mosaicism of other F_1_ genotypes was again absent except in crosses involving the 2093A line where it was present only in *kmo^sgRNA^* individuals. Interestingly, however, here it occurred both where Cas9 had been deposited by the male (14.0% F_1_ mosaic) and female (20.4% F_1_ mosaic) parents.

**Table 3.**
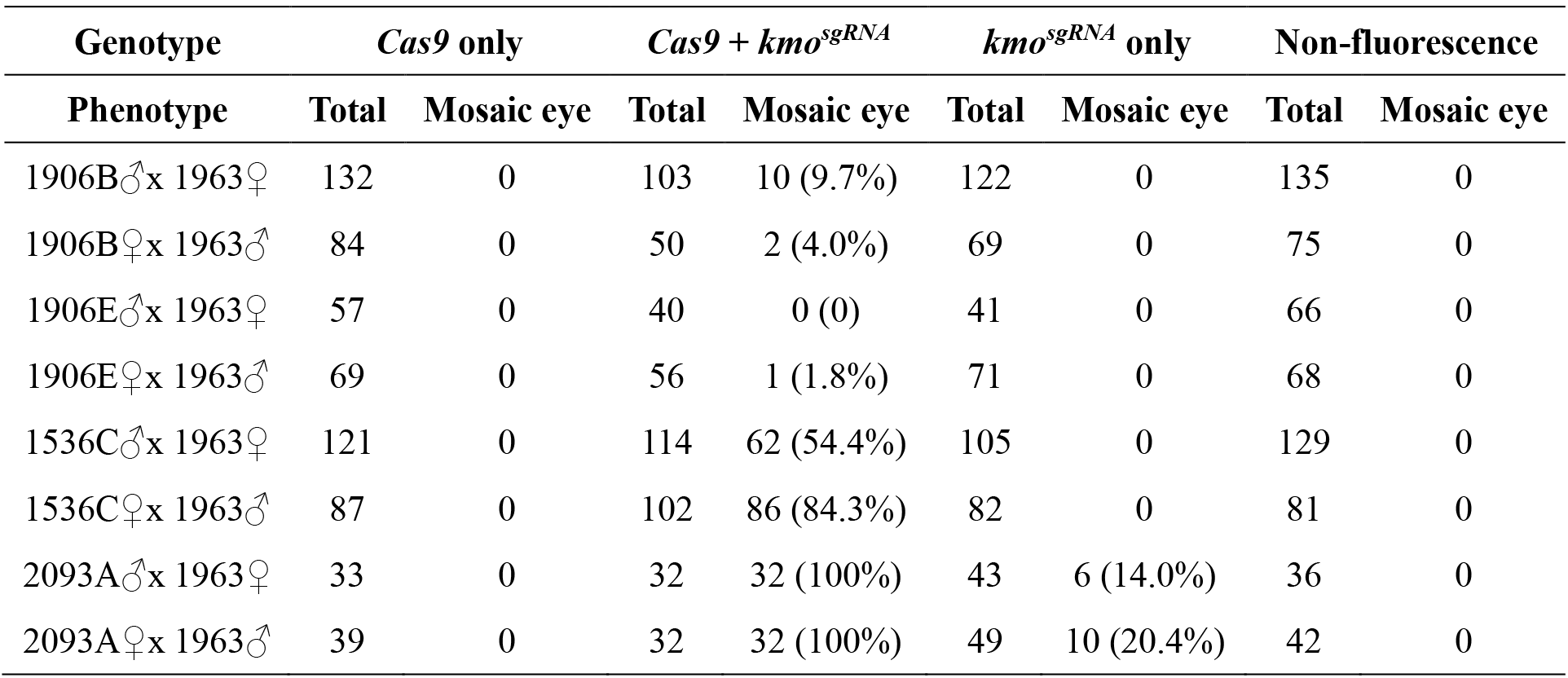
Individuals and percentages of different genotypes and phenotypes shown in F_1_s generated from *Pxkmo* homing crosses.

| **Genotype** | ***Cas9* only** |  | ***Cas9* + *kmo*<sup>sgRNA</sup>** |  | ***kmo*<sup>sgRNA</sup> only** |  | **Non-fluorescence** |  |
| --- | --- | --- | --- | --- | --- | --- | --- | --- |
| **Phenotype** | **Total** | **Mosaic eye** | **Total** | **Mosaic eye** | **Total** | **Mosaic eye** | **Total** | **Mosaic eye** |
| 1906B♂x 1963♀ | 132 | 0 | 103 | 10 (9.7%) | 122 | 0 | 135 | 0 |
| 1906B♀x 1963♂ | 84 | 0 | 50 | 2 (4.0%) | 69 | 0 | 75 | 0 |
| 1906E♂x 1963♀ | 57 | 0 | 40 | 0 (0) | 41 | 0 | 66 | 0 |
| 1906E♀x 1963♂ | 69 | 0 | 56 | 1 (1.8%) | 71 | 0 | 68 | 0 |
| 1536C♂x 1963♀ | 121 | 0 | 114 | 62 (54.4%) | 105 | 0 | 129 | 0 |
| 1536C♀x 1963♂ | 87 | 0 | 102 | 86 (84.3%) | 82 | 0 | 81 | 0 |
| 2093A♂x 1963♀ | 33 | 0 | 32 | 32 (100%) | 43 | 6 (14.0%) | 36 | 0 |
| 2093A♀x 1963♂ | 39 | 0 | 32 | 32 (100%) | 49 | 10 (20.4%) | 42 | 0 |

A random subset of mosaic transheterozygotes were collected for both 1963C (<u>*Pxkmo*</u>) and 1619P15 (<u>*Pxyellow*</u>) crosses and editing events confirmed with a T7E1 mutagenesis assay (Figure S4).

### Germline activity of CRISPR/Cas9 system (F_2_ analysis)

F_1_ transheterozygotes derived from the above experiments were reciprocally mated to relevant knockout lines (*Pxyellow*−/− for F_1_s carrying 1619P15 and *Pxkmo*−/− for F_1_s carrying 1963C/1962A), and their F_2_ offspring were then scored for fluorescence and knockout phenotypes (Figure 3). Under Mendelian inheritance, 50% of these F_2_ would be expected to carry the relevant sgRNA-expressing transgene with significant deviation above this taken to be an indication of inheritance bias caused by gene drive activity. For the control crosses, heterozygous Cas9 and sgRNA lines were independently, reciprocally, mated to the two knockout lines with F_2_ data collected as above (Figure S5).

**Figure 3.**
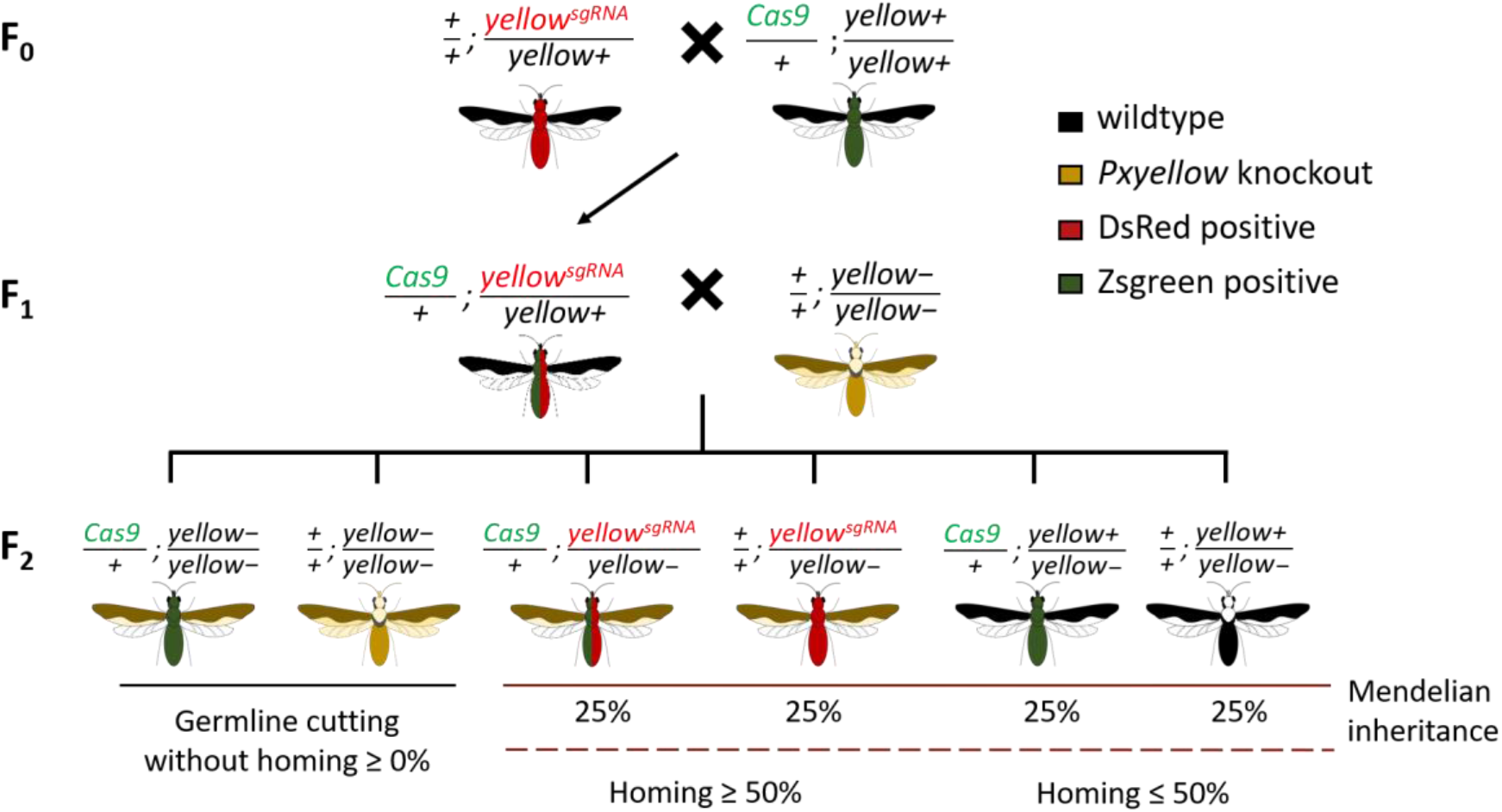
Crossing scheme for the experiment. Taking *Pxyellow* cross group for example, based on mendelian inheritance, only two pigmentation phenotypes as well as four genotypes will be observed in the F_2_ generation: 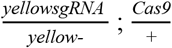 (yellow pigmentation, DsRed positive, Zsgreen positive), 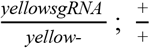 (yellow pigmentation, DsRed positive, Zsgreen negative), 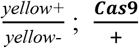 (wildtype pigmentation, DsRed negative, Zsgreen positive), 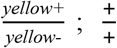 (wildtype pigmentation, DsRed negative, Zsgreen negative), each representing 25% of the progeny, thus making the *yellowsgRNA* (homing element) individuals = 50%. However, if germline cleavage and subsequent NHEJ-based repair occurred, another two types: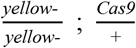 (yellow pigmentation, DsRed negative, Zsgreen positive),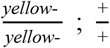 (yellow pigmentation, DsRed negative, Zsgreen negative) will occur. If homing occurs in F_1_ germlines, we would expect a bias in favour of individuals inheriting the *yellow^sgRNA^* allele in the F_2_ (>50%) at the expense of those not inheriting this element (<50%). As for targeting *Pxkmo*, in contrast to dark compound eyes in wildtype and heterozygous adults, homozygous null mutation of *Pxkmo* resulted in yellow eyes.

Results showed no significant deviation from 50% inheritance for any of the three sgRNA-expressing transgenes assessed at a significance threshold of *P*≤ 0.05 (Figure 4A), indicating that any gene drive activity, if present, was too low to detect. Interestingly, however, the observation of some WT and Cas9-only individuals with full *Pxkmo* or *Pxyellow* knockout phenotypes in the F_2_ progeny indicated that germline expression and activity of Cas9 was nonetheless taking place (Figure 4B). These full knockout phenotypes could either have derived from cleavage of a parental WT chromosome in the F_1_ germline, or cutting in the F_2_ following deposition of Cas9 by the F_1_. Given that these individuals displayed full knockout phenotypes (complete white eyes or yellow bodies) rather than the mosaic phenotypes observed in the F_1_ analysis (where no such full knockout phenotypes were observed), we believe the former (germline cutting) hypothesis to be the more likely. It cannot be ruled out that very early cutting in the F_2_ embryo may have contributed to this observation, however. Regarding 1619P15 (Figure 4B-a), this activity was present in crosses involving the 1536 and 2093 but not 1906 lines. Comparing putative germline cleavage rates by sex of transheterozygous F_1_ parent between Cas9 lines, we found that 1536C and 2093A lines showed the highest average cleavage when assessing F_1_ male parents (6.0% and 7.7%, respectively) whereas 1536C exhibited the highest average germline cutting when assessing F_1_ female parents (2.9%). When comparing between sexes within each Cas9 line, the germline Cas9 activity in male transheterozygous parents was significantly higher than in female parents for both 1536C and 2093C crosses. Other comparisons were non-significant.

**Figure 4.**
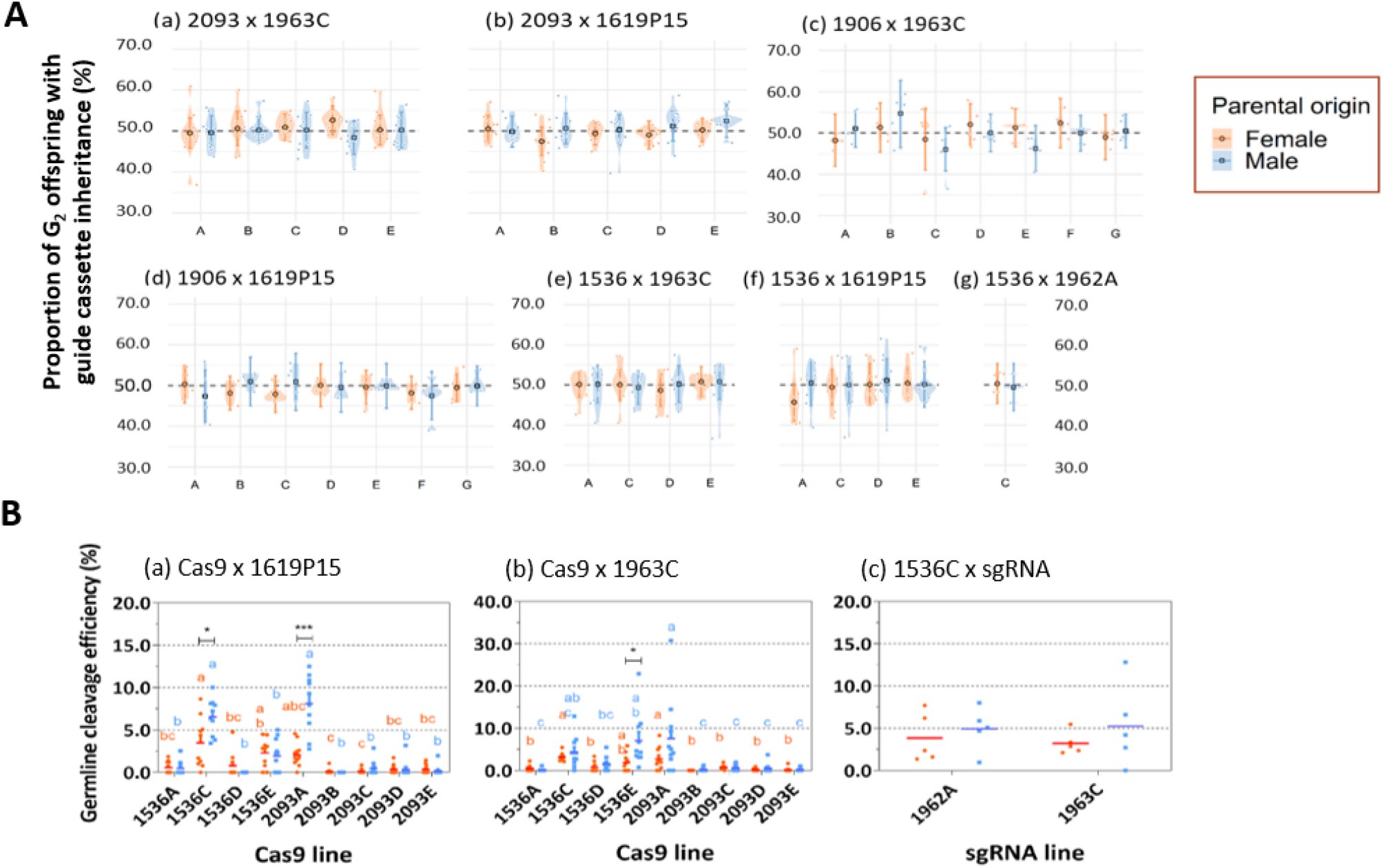
Assessment of the inheritance of sgRNA cassettes and the germline cleavage efficiency. A: Proportion of F_2_ individuals inheriting the relevant homing element. Offspring come from either a male (blue) or female (orange) Cas9-bearing F_1_ parent. Due to non-significant effect of ‘grandparental’ (F_0_) sex on the model, this factor has been collapsed in the data shown. Central tendency and error bars represent estimated mean proportions and associated approximate 95% confidence intervals for that treatment. Shaded areas (violin plots) represent the density distribution of the raw data. B: Germline cleavage efficiency (%) = no. phenotypic mutants (e.g. yellow eyes) / (no. phenotypic mutants + phenotypic wildtypes (e.g. black eyes)). Offspring come from either a male (blue) or female (orange) Cas9-bearing F_1_ parent Data are plotted as means overlaid on raw data. For comparison of lines within the same sex, those which share a letter above the plotted data (e.g. ab and bc) are not significantly different from each other. For comparisons between sex, within each line, “*” shows the significant difference between subgroups. *: *P* < 0.05. **: *P* < 0.01. ***: *P* < 0.001.

Similarly for crosses involving 1963C (Figure 4B-b), putative germline cutting events were detected exclusively in F_1_ WT and Cas9-only individuals from 1536 and 2093 crosses. Specifically, 2093A showed the highest germline editing levels, independent of whether assessing male or female F_1_ transheterozygous parent (male F_1_ = 7.6%, female F_1_ = 2.8%. It was also noted that male F_1_ parents showed higher germline cleavage rates than females in the 1536E crosses. To compare the cleavage efficiency induced by different *PxU6* promoters, we also conducted 1536C x 1962A crosses. Although null-mutants derived from germline cleavage were also found in F2s, no significant differences in frequency of these events were observed between 1962A and 1963C crosses involving 1536C (Figure 4B-c).

A random selection of WT F_2_ knockout mutants were individually sequenced at the relevant target site. Multiple indel alleles were observed separate from those previously established as occurring in each knockout line^26^ (Figure S6).

Finally, gonads and carcasses of WT, 1536C, 2093A and 2093D heterozygotes were dissected for comparing Cas9 protein expression. At a broad scale, it was noted that Cas9 protein levels generally appeared higher in carcasses than gonads given equal amounts of total protein loaded (35μg), in all three transgenic lines, while no Cas9 was expressed in WT (Figure S7), a result in line with Cas9 transcription patterns detected by RT-PCR (Figure S1), However, unlike Cas9 mRNA patterns, Cas9 protein was not observed in 2093D gonads. This result concurs with the observation of both somatic and germline Cas9 activity (cutting) in the 1536C and 2093A lines but no germline activity in the 2093D line, strengthening the argument that cutting events detected in the F_2_ generation resulted from Cas9/sgRNA activity in the F_1_ germline.

## Discussion

Previous studies integrating CRISPR/Cas9 components into lepidopterans are limited to the highly domesticated model species *Bombyx mori* (silkworm). There, using the IE1 promoter to drive Cas9 resulted in somatic editing of transheterozygotes^39–41^, while using the *B. mori nanosP* promoter resulted in both somatic mosaicism of transheterozygotes^30,42–53^ and some germline activity^54–56^. However, these studies provide no rates for these activities and due to their experimental design, it is often difficult to disentangle the effects of germline edits from somatic mosaicism. Here, we expand the use of integrated CRISPR/Cas9 components to a free-living agricultural pest of global importance, using an experimental design which allowed us to quantify and clarify different editing activities, and also assess the possibility of homing-based gene drive. Using three different endogenous promoters, which were putatively germline-specific, we developed Cas9 expressing DBM lines. Testing these lines, we were able to demonstrate all three forms of Cas9 nuclease activity – *somatic expression*, resulting in mosaicism of F_1_ transheterozygotes: *trans-generational deposition*, resulting in F_1_ *target^sgRNA^*-only mosaics, and *within germline expression*, resulting in F_2_ non-fluorescent (WT) and Cas9-only target gene full-knockouts. Our results indicate that, in general, *Pxvasa-Cas9* and *PxnanosP-Cas9* drove higher rates of Cas9 activity than *Pxmeiw68-Cas9*.

We observed that somatic Cas9 expression from somatic activity of the *vasa* and *nanosP* promoters^14–16,57^ was much enhanced relative to native *vasa* and *nanos*P, in all Cas9 lines. This may be an interaction with the Hr5 enhancer used to aid expression of the fluorophore marker in our constructs. Although *nanosP* promoter activity has been reported as more germline-restricted than *vasa* in *Drosophila*^57,58^, these two promoters both achieved high levels of somatic Cas9 expression in DBM, resulting in 100% observable mosaicism in multiple lines. In order to reduce this somatic overexpression, future studies could employ alternative regulatory elements without long distance enhancers to drive marker expression, such as the 3xP3 or OpIE2 promoters, both of which have been characterised in lepidopteran transgenesis^32^. Nonetheless, ‘leaky’ somatic expression of Cas9 may be a significant problem in constructing population-suppression gene drives targeting fertility/viability genes, due to the generation of resistant alleles and the high fitness cost in drive heterozygotes^59,60^.

Both maternal and paternal deposition of Cas9 occurred in *PxnanosP-Cas9*, although maternal Cas9 was more abundant, judging by mosaicism of *target^sgRNA^* -only F_1_ progeny. Our results contrast with a similar study in *Drosophila melanogaster*, where maternal but not paternal deposition of Cas9 was evident^61^. Similar deposition, however, was not observed in *Pxvasa-Cas9* lines although such carryover occurs at high levels in *vasa-Cas9 Anopheles gambiae*^59^ and *vasa-Cas9 D. melanogaster*^61^ transgenics, and Cas9 protein was clearly detected in the ovaries and testes of a *Pxvasa-Cas9* line here. It is possible that deposition of Cas9 protein or mRNA occurred in *Pxvasa-Cas9* lines but not at a level high enough to be observed in the relatively small areas (eyes and dorsal stripes) where mosaicism could be detected.

Germline activity of Cas9 was confirmed by the hypothesised inheritance of null *Pxkmo*/*Pxyellow* alleles by F_2_ non-cas9 (WT and *target^sgRNA^*) individuals. However, compared to studies *in D. melanogaster*^60,61^ where both *vasa* and *nanos* promoters have been tested, rates of germline cutting were relatively low, with a maximum mean of 8.1±1.0% of wild-type chromosomes mutated. This was surprising as the high rate of somatic mosaicism observed suggested the expressed Cas9 protein and Pol III sgRNAs were highly capable of acting at the target site. A possible explanation could be either insufficient Cas9 expression/translation or sgRNA expression from the employed promoters within germline cells. While RT-PCR and western blot analysis of gonads from Cas9 lines confirmed the presence of Cas9 mRNA/protein, we cannot exclude the possibility that this may have largely represented expression within the somatic cells of gonadal tissues. As in studies assessing CRISPR-based split-drive in *Aedes aegypti*^17^, we observed that Cas9 lines generated with the same construct varied in Cas9 expression (RT-PCR and Western blot) and subsequent germline cleavage rates, likely due to the “positional effect” of semi-random *piggyBac* insertion. Future studies may seek to employ a site-specific approach once suitable ‘safe-harbour’ sites have been identified, or an ‘integral’ gene drive approach where endogenous regulatory elements can be hijacked for canonical Cas9 expression^62^. Interestingly, when comparing between sexes within each cross, significant differences (where they occurred) showed a higher level of cleavage in male rather than female germlines, whereas the opposite trend would have been expected given the endogenous expression patterns of *Pxvasa* and *PxnanosP*.

Germline editing and conversion rates can vary according to the locus being targeted by the homing element, likely affected by chromosome/chromatin structure, sgRNA efficiency or sequence diversity in the population^15,60^. As such, we constructed sgRNA expressing lines by site-specifically inserting guide cassettes into two loci – *Pxkmo* and *Pxyellow*. To our knowledge, this is the first report of HDR-based long fragment knock-in using CRISPR/Cas9 in the Lepidoptera, although site-specific integrations using the more complex MMEJ-based PITCH technologies and TALEN-based integration have been demonstrated in *Bombyx mori*^45,63,64^. However, knock-in efficiency was relatively low (0.4-0.7%), compared with highly efficient knockout in DBM (57.1%)^26^. Integration rates were also lower than comparable reports in dipterans^65–67^. Interestingly, our results showed a large number of non-canonical repair events, with only two of the seven generated homing elements lines showing a precise, perfect integration. This runs contrary to experience in other insect species published^17,57,65,68^, and also our extensive experience in mosquitos, where off-target integration may occur relatively often, but those constructs integrated at the correct location are almost always integrated perfectly. The reasons for this difference are unknown but it is noted that previous efforts to generate Cas9-mediated insertions using plasmid templates in *B. mori* have failed, while similar TALEN-based integrations are relatively efficient^45^. Therefore, it may be that some part of the DSB repair process in lepidopterans differs from other insects studied so far, reducing the efficiency of donor single strand invasion following the blunt ends breaks created by Cas9, but not the overhanging breaks created by TALENs. Other genome engineering tools (e.g. TALEN^45,63^, TAL-PITCH and CRIS-PITCH^64^) could be employed to increase the integration efficiency of CRISPR/Cas9 split-drive components in DBM, although for this purpose the integration site mediated by these technologies would need to fortuitously coincide with the chosen sgRNA cleavage site. Additionally, we successfully utilised Cas9-based targeted sequencing of the integration junction to resolve the location of one of these homing element lines (1619P15). To our knowledge this is the first published use of such a system and may be of benefit to other researchers seeking to resolve transgene integration junctions that are similarly difficult to resolve using conventional PCR-based techniques.

Despite testing three putatively germline-active Pol II promoters and six U6 Pol III promoters at two independent loci, we were unable to demonstrate significant gene drive. One explanation for this could include insufficient expression of Cas9 mRNA or its translation in tissues and at times required for a ‘homing’ reaction to occur. The simultaneous presence of Cas9 protein and sgRNA in a germline-specific time window is critical for homing, especially in early meiosis when the germ cell is in a recombination-orientated state and homologous chromosomes are in close enough proximity to act as HDR templates^61,69^. We were able to demonstrate here that Cas9 protein was present in gonadal tissues of both sexes and that it was able to mediate heritable germline DNA breakage events after complexing with sgRNAs, however, the timing of these events may have presented before or after this narrow window, and/or the expression levels might be too low in germ-cells to achieve distinguishable biases in inheritance, as has been similarly reported recently in mice^70^. As an additional hurdle, the imperfect repair of the 5’ homology arm/genome interface in 1619P15 and 1962A may have reduced the ability of these integrations to act as repair templates, although in 1619P15 there remained at least 257 bp of homology arm sequence on this 5’ end. This however did not apply to 1963C, which was a precise integration. Finally, we are unable to discount the possibility that the six known U6 promoters in DBM do not function efficiently in the germline. We recommend additional germline-specific regulatory components to be tested in DBM to increase germline cleavage and conversion efficiency.

## Conclusion

In summary, this study has identified and characterized endogenous genetic components (i.e., germline active Pol II and Pol III promoters) and completed the first CRISPR/Cas9-mediated site-specific knock-in of large fragments in Lepidoptera. With these components, we built and tested the first split-drive system in a lepidopteran, achieving a very high level of somatic editing. Although we did not see significant homing, Cas9-mediated germline cleavage as well as maternal and paternal Cas9 deposition was observed. Our results provide valuable experience, paving the way for future construction of gene drive-based genetic control strategies in the global pest, DBM, or other lepidopterans.

