## Supplementary material for "Towards CRISPR/Cas9-based gene drive in the diamondback moth *Plutella xylostella*": Figure S

### Supplementary data

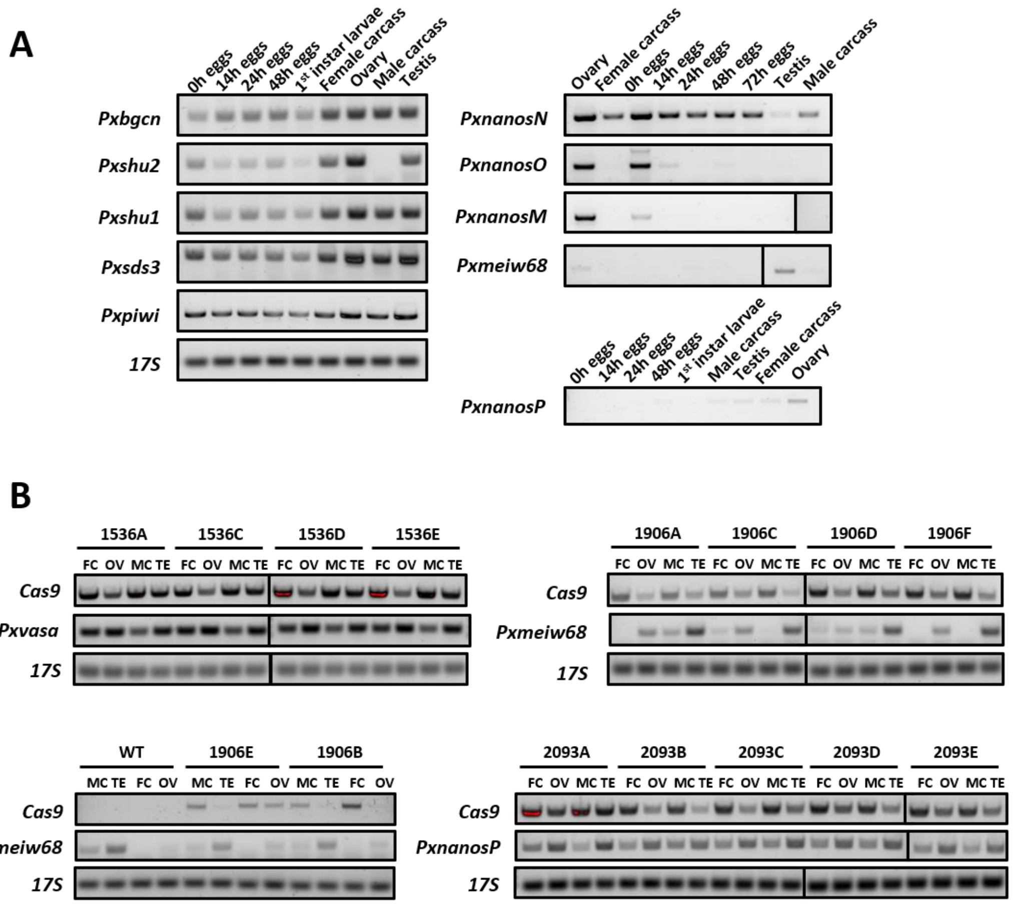


**Figure S1 Expression profiling of endogenous germline candidate genes in wildtypes (A) and both endogenous and exogenous genes in transgenic lines (B).** *17S* is broadly expressed and used as a loading control. Tissue samples are all dissected from adults (carcass indicates moth tissues excluding dissected gonads). FC: female carcass. MC male carcass. OV: ovary. TE: testis.

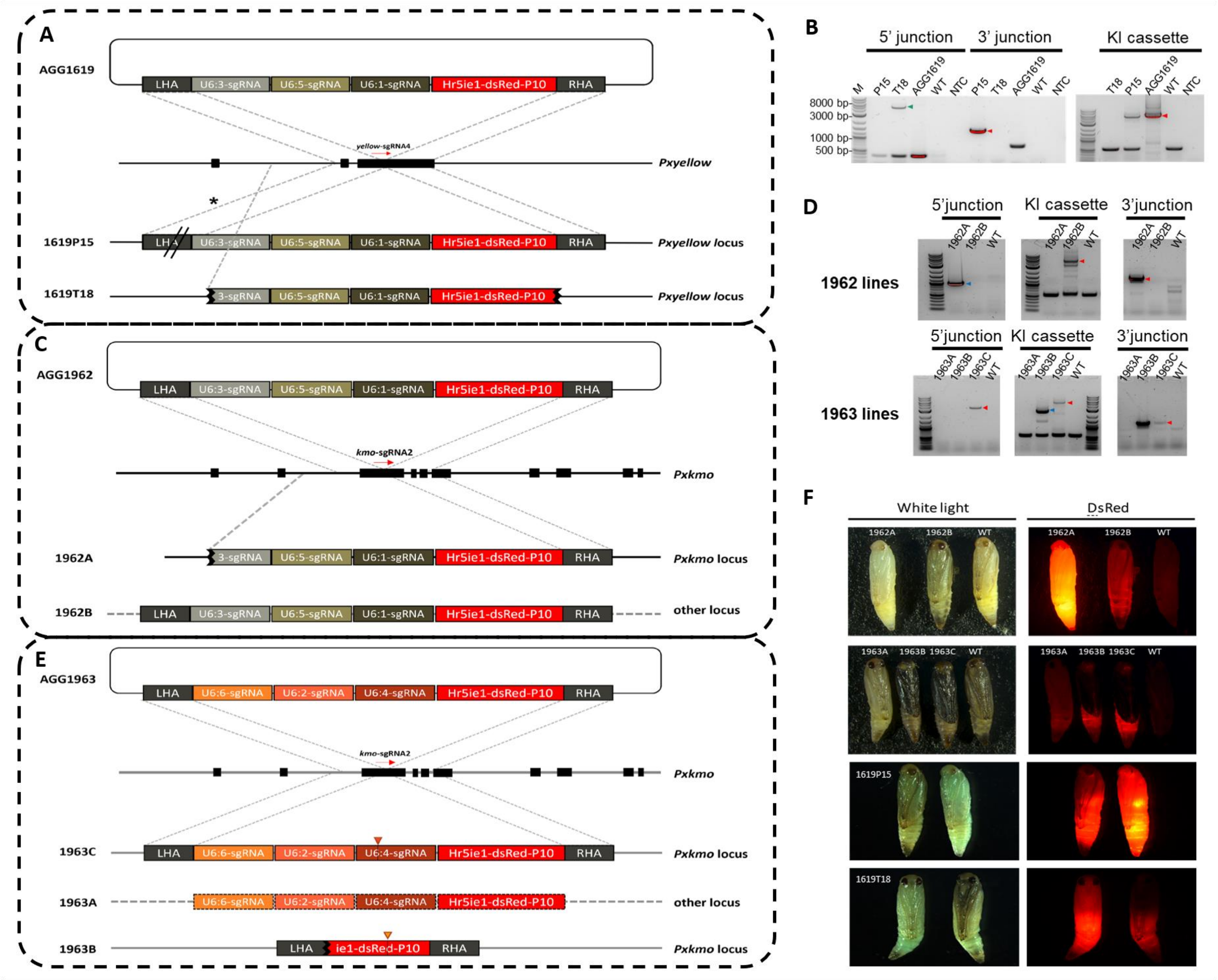


**Figure S2 CRISPR-based integration of sgRNA cassette into *Pxyellow* and *Pxkmo* loci.** A: Designed knock-in donor plasmid AGG1619 and the deduced genetic structure of two transgenic lines (1619P15 and 1619T18). Based on the PCR result in Figure S2-B, unconfirmed 5' junction of 1619P15 is marked with "//". Sequencing the largest PCR product which could be amplified downstream of this junction confirmed the presence of at least 257bp of the left homology arm (LHA) but we could not produce an amplicon extending further than this upstream - either into the remainder of the homology arm, nor the 5' genomic flanking region. However, this junction was subsequenctly succesfully confrmed using Cas9-based targeted sequencing of the integration (Oxford Nanopore Technologies). The subsequently confirmed junction is marked with *. The 5' fragment, including partial PxU6:3 promoter and the left homology arm, is lost in 1619T18 line, of which 3' junction is likely also deficient. B: PCR verification of 5' and 3' junctions as well as knock-in cassette in 1619 lines. Blue arrows indicate bands smaller than expected, while the red ones point out bands of correct sizes. Amplicons were sequenced to confirm. C and E: Knock-in donor plasmid AGG1962 and AGG1963 as well as the deduced genetic structure of transgenic lines (1962A, 1962B, 1963A, 1963B and 1963C). The knock-in cassettes in 1962B and 1963A were likely integrated into off-target loci, while 1962A line lost partial 5' sequence (left homology arm and part of *PxU6:3* promoter). Besides, 1963B showed a large deletion in the donor cassette (inverted yellow triangle), while the 1963C line displayed a 2 bp deletion inside the *PxU6:4* promoter (inverted orange triangle). D: PCR verification of 5' and 3' junctions as well as knock-in cassette in 1962 and 1963 lines. E: knock-in donor plasmid AGG1963 and the deduced genetic structure of three transgenic lines (1963A, 1963B and 1963C). F: The transgenic lines show strong fluorescence under the DsRed filter. In A, C and E, location of the target sequence for each respective sgRNA is shown by a horizontal red arrow and labeled.

AGCCTCCTATTGCTTACTGTTAGGTATGGGACTATAACTTCGTATAATGTATGCTATACGA
GAAGTTATGTCCCGGCATGAAGAGTTGACGTGAGTGAACCGCCAAGATTTGTTCTGCT
CCCAGGAGTAGGCGATGAGTCCTATCCGGGTTCGTCCGGAAAAGTATGCAAGGTGTCT
TCACAGCTTTTTGCGCTTCATCGGAAGCAATGTTAGCGATGAAAGTGGTGGACGTGTC
TTCAGGGCGGAAGACGTATCTCCCGGATACGACGGTCAGTTTTAAGATCATAGACGTTA
AGGGCATAGGACAGGGGTTAGTAACATTGGGATCACAAAACAAACAAAACATTTGTTT
ATAAATAATTAACCTTGTAAATAATGAAATTGGCTAATCTACGAATTGAAGAAGGCAGAT
GTGATATTGCTCGATTCGCTGCGTCATTCTATCGTCGTATTACAAACAGTAAAATTAGCT
TTTATTTCGGCATGGAAATCATTTATAAACTATGAAAAGTAAATTATTTTATAGCAAATAA
ACTGTCAGAATTGATACTAACCATAACCATAAGTGCCAACATCAACGCAGAGACTGGT
CGCATTTGTCGGCTTTGATTCTGTAGACGGTGTTCAGTCCATTCTGACAGTTGCCGATTT
TCATTGCCCTCCCCAGCTGGGGTGGGGGGATCAACTTGGGGGATGGCTCGTAAGGGGC
ATCGAGGGGATGTAGTTCAATGTGGCAGGATACCTGGAATAGATGAAAAAATAAAGTTT
AGCAACTCTTTCAAATTAATTTTATTTAAGGAGAACCTAGACCTGAAAATATAAAGAGA
TTAAATTTAACATAAAGTTCAACTGTTTAGATGGTGGTTTGGTGGTCCACCATTGCCAG
AGAAGTATCTATCTTAATATATATATATATATATATAATATATTTATTTATTTAATTACTTTATT
GCTGAATATAAAATAAATGTACAAAGATGGGCTTAATAACTAGAGCATATATATATAATAT
ATTATATAATGTATTATATATATAATATATATAAACTA

**Figure S3 Resolved 5’ junction of 1619P15 integration using Cas9-based targeted sequencing.** Red lettering: Terminus of Pxy U6.3. Green lettering: attB site. Blue lettering: Homology arm included in construct. Orange lettering: genomic flanking sequence

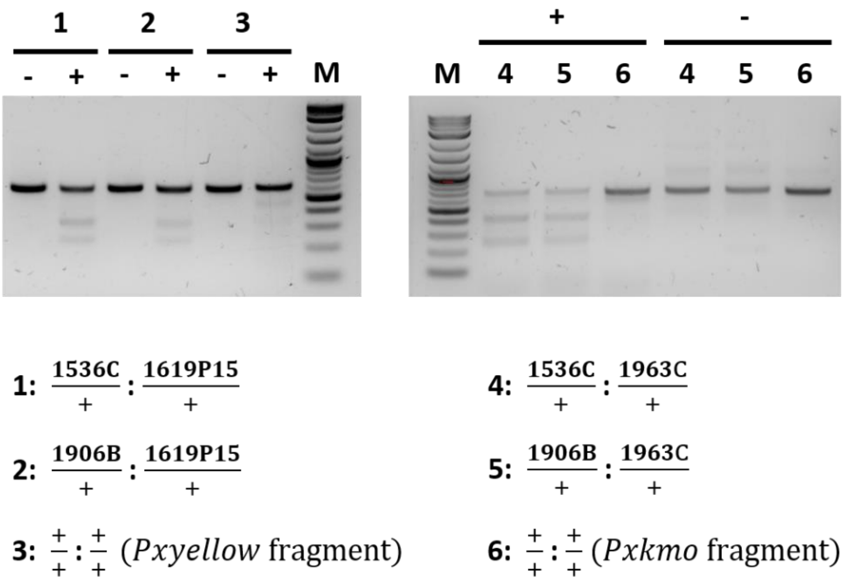


**Figure S4 T7E1 of $G_1$ transheterozygous pupae showing somatic mosaicism.** Mutations in *Pxyellow* and *Pxkmo* target sites were confirmed by two expected bands derived from T7 endonuclease cleavage. In gel pictures, "+" and "-" indicate with and without T7 endonuclease treatment. In the figure legend, "+" means wildtype allele.

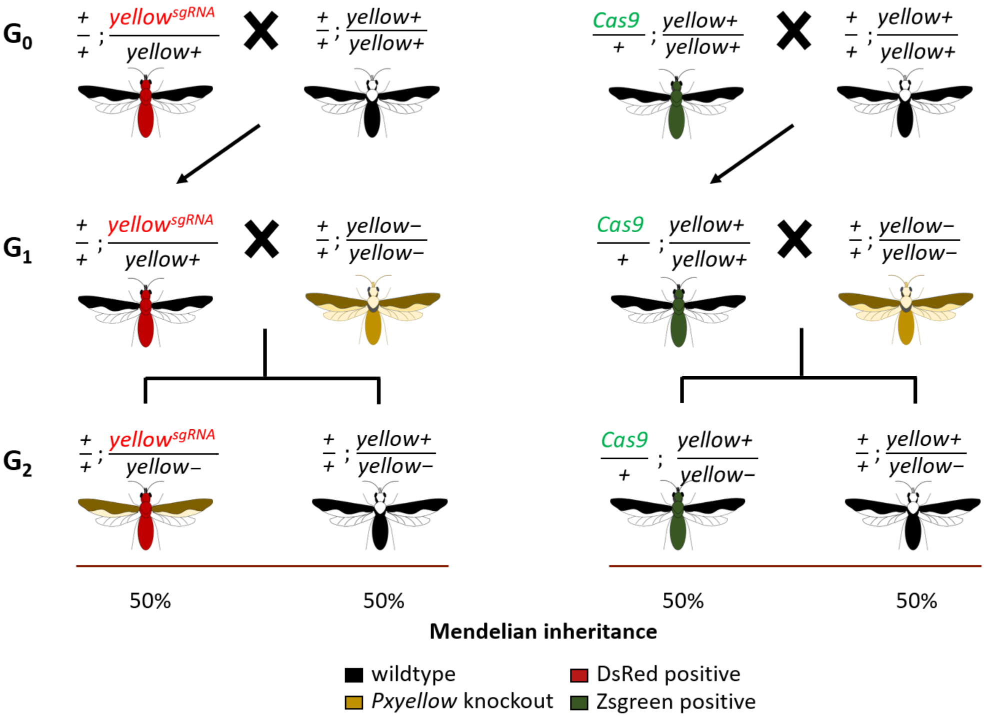


**Figure S5 Control cross scheme.** Crosses are set for excluding the possibility of lines contamination, as well as providing data as intercept in the statistical analysis of homing element germline inheritance in Figure 3.

**A**

| Line | Mutation | Sequence |
|---|---|---|
| WT | - | CACTCACAACTTCTTCATGCCTGACCCTCTTGTTGGTGAATTCAACATCGCCGGTCTGACCTTC |
| *Pxyellow*-KO | -192 bp, +12 bp | TCGTATTCACGCCGTA / -192 bp / CCTCTTGTTGGTGAATTCAACATCGCCGGTCTGACCTTC |
| 1536E | -8 bp | CACTCACAACTTCTTCATGCCTGAC--------TGGTGAATTCAACATCGCCGGTCTGACCTTC |
| | -2 bp | CACTCACAACTTCTTCATGCCTGA--CTCTTGTTGGTGAATTCAACATCGCCGGTCTGACCTTC |
| 1536C | -1 bp | CACTCACAACTTCTTCATGCCTGA-CCTCTTGTTGGTGAATTCAACATCGCCGGTCTGACCTTC |
|  | -9 bp | CACTCACAACTTCTTCATGCCTGAC---------GGTGAATTCAACATCGCCGGTCTGACCTTC |
| 1536A | +13 bp | CACTCACAACTTCTTCATGCCTGACTTCATGCCTTCTTCCTCTTGTTGGTGAATTCAACATCGCC |
|  | -12 bp | CACTCACAACTTC------------CCTCTTGTTGGTGAATTCAACATCGCCGGTCTGACCTTC |
| 1536D | -1 bp | CACTCACAACTTCTTCATGCCTGA-CCTCTTGTTGGTGAATTCAACATCGCCGGTCTGACCTTC |
| 2093C | -15 bp | CACTCACAACTTCTTCATGCCTGAC---------------TTCAACATCGCCGGTCTGACCTTC |
|  | -12 bp | CACTCACAACTTCTTCATGCCTGAC------------GAATTCAACATCGCCGGTCTGACCTTC |
| 2093A | -4 bp | CACTCACAACTTCTTCATGCCTGA----CTTGTTGGTGAATTCAACATCGCCGGTCTGACCTTC |
| | -356 bp, +16 bp | TCCCCACCACTTTCATCGTCCTGACCGCGTCAG / -356 bp /CCGCGGCCCCGACACCCACAC |
| 2093C | -10 bp | CACTCACAACTTCTTCATGCCTGACC----------TGAATTCAACATCGCCGGTCTGACCTTC |
|  | -12 bp | CACTCACAACTTCTTCATGCCTGAC------------GAATTCAACATCGCCGGTCTGACCTTC |
| 2093E | -8 bp | CACTCACAACTTCTTCATGCCTGAC--------TGGTGAATTCAACATCGCCGGTCTGACCTTC |
|  | -7 bp | CACTCACAACTTCTTCATGCCTGAC-------TTGGTGAATTCAACATCGCCGGTCTGACCTTC |
| 2093D | -5 bp | CACTCACAACTTCTTCATGCCTGAC-----TGTTGGTGAATTCAACATCGCCGGTCTGACCTTC |

**B**

| Line | Mutation | Sequence |
|---|---|---|
| WT | - | ACATACGGAACACACCACAGGTCAGAGGGCGGTCC/-120bp-/TCTACGTATGAGATACCGTACGATGCGAGGACCAATCAG |
| *Pxkmo*-KO | -10 bp | ACATACGGAACACACCACAGG-----GGGCGGTCC/-120bp-/TCTACGTATGAGATACCG-----TGCGAGGACCAATCAG |
| 1536E | -4 bp | ACATACGGAACACACCACAGGTCAGAGGGCGGTCC/-120bp-/TCTACGTATGAGATACCGT----TGCGAGGACCAATCAG |
|  | -7 bp | ACATACGGAACACACCACAGGTCAGAGGGCGGTCC/-120bp-/TCTACGTATGAGATAC-------TGCGAGGACCAATCAG |
| 1536C | -1, +5 bp | ACATACGGAACACACCACAGGTCAGAGGGCGGTCC/-120bp-/TCTACGTATGAGATACCGTACCATAT-ATGCGAGGACCA |
| | -2 bp | ACATACGGAACACACCACAGGTCAGAGGGCGGTCC/-120bp-/TCTACGTATGAGATACCGTAC--TGCGAGGACCAATCAG |
| 1536D | -5 bp | ACATACGGAACACACCACAGGTCAGAGGGCGGTCC/-120bp-/TCTACGTATGAGATACCG-----TGCGAGGACCAATCAG |
| 1536A | +7 bp | ACATACGGAACACACCACAGGTCAGAGGGCGGTCC/-120bp-/TCTACGTATGAGATACCGTACGATGGAGGATGCGAGGAC |
| 2093A | -1 bp | ACATACGGAACACACCACAGGTCAGAGGGCGGTCC/-120bp-/TCTACGTATGAGATACCGTACGA-GCGAGGACCAATCAG |
|  | -8 bp | ACATACGGAACACACCACAGGTCAGAGGGCGGTCC/-120bp-/TCTACGTATGAGATACCGT--------AGGACCAATCAG |
| 2093C | +1 bp | ACATACGGAACACACCACAGGTCAGAGGGCGGTCC/-120bp-/TCTACGTATGAGATACCGTACGATTGCGAGGACCAATCAG |
| 2093E | -2 bp | ACATACGGAACACACCACAGGTCAGAGGGCGGTCC/-120bp-/TCTACGTATGAGATACCGTAC--TGCGAGGACCAATCAG |

**Figure S6 Mutant alleles generated via germline cleavage and subsequent NHEJ-mediated repair in *Pxyellow* (A) and *Pxkmo* (B) loci.** The left column shows corresponding Cas9 cross groups where these $F_2$ mutations originated from. *Pxyellow*-KO and *Pxkmo*-KO respectively represent the knockout line used in $F_1$ crosses. Indel genotypes are listed in the middle column, while the sequence details are shown on the right. sgRNA target region is marked in red while the PAM site is underlined. Blue letters signify inserted bases, and dashed lines indicate deletion. Partial fragments are omitted and shown in numbers, because the sequences are too long to be presented in the figure.

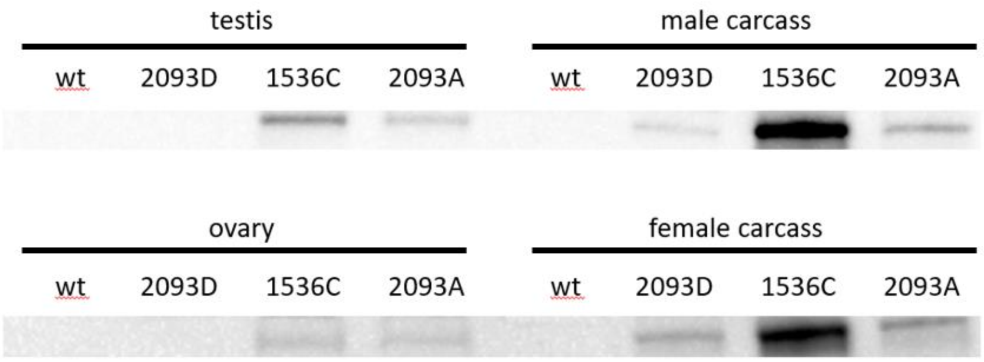


**Figure S7 Cas9 protein expression in gonads and carcasses of WT, 1536C, 2093A and 2093D lines.** Equal amounts of protein were loaded for each sample.

**Table S1 Primers used this study**

| Primer ID | Primer sequence (5' to 3') | Target gene fragment | Aim |
|---|---|---|---|
| LA1126 | AAGAACTCGCGTGAACGTATGAAG | *Cas9* | RT-PCR |
| LA3491 | TCACTTGTGGCATCGACAGCACC | | |
| LA309 | CAAGCCTCTTCGTAACAAGATCG | *17S* | |
| LA310 | CAGCTTGATGGAGATACCACGC | | |
| LA4444 | AGTGACCCACATATTCGTGG | *PxBGCN* | |
| LA4445 | GTTCTTCTGACGGCTGCTTC | | |
| LA4460 | AACGGCATAGCATGGAGCTG | *PxSDS3* | |
| LA4463 | TCGTCAGTGATGGCGTGTATG | | |
| LA4453 | ACCACGGATCAAGCCTAAAG | *Pxshu1* | |
| LA4455 | TCTCGTTGCAAGCAGTGCAAGC | | |
| LA4449 | AGAGTTGGTCCCGTACGATG | *Pxshu2* | |
| LA4451 | TGCGTTAGATGCAGCTACAG | | |
| LA3198 | AGGCAACTTCTGTCTCGGACTGACGAAAC | *Pxmeiw68* | |
| LA3199 | ACTATGATGTTGAATCGAGTTCTGTCTTCAC TGTTAG | | |
| LA4456 | ACCAGATCAGGATCCACGAC | *Pxpiwi* | |
| LA4458 | ACCCTTGATCTTCACCAACG | | |
| LA3213 | AATGATAATTACTACAGTGACTATGGAAGT GCCAGTTC | *PxnanosO* | |
| LA3216 | TCGCCGTTGTTCTTGCAGAATCTGCATTC | | |
| LA3201 | GGAGTCTCTGGCCTATAATTTTGCTATGGAA GATAC | *PxnanosN* | |
| LA3204 | TTGGAGTCGGTCTCGTCGAAGGCCATC | | |
| LA3205 | ATAACCTAACAACACTGTTCAAGGACACAC TGTCAAC | *PxnanosM* | |
| LA3207 | AATACCACGGCGGGGACTCGCCGTTG | | |
| LA5967 | CAACCATGCAACCCTGACACTAGGA | *PxnanosP* | |
| LA5968 | TTAGTACCTCCGTTTCCCGTTACTCTTG | | |
| LA4722 | CTGCATGGTGATCCACCTGGTCTCATAG | *Pxmeiw68* promoter | Assembly of Cas9 constructs |
| LA4721 | CATTTGACGAAGATCTTTCGGCATCTTAAA | | |

| | | | |
|---|---|---|---|
| | C | | |
| LA4703 | ACTACGAAAATAAATACAAGTTAC | *Pxmeiw68* 3'UTR | |
| LA4704 | TCAATGTATCTTAACGCGAGTTAATTTCAGAACATCTACACTG | | |
| LA6008 | CGATTCGAGTTAACGGCCGGGCAAAGCCCACGAATTTGCACT | *PxnanosP* promoter | |
| LA6009 | TCGTGGTCCTTATAGTCCATTTTCATCTGAAATGACACAATTTTAACACG | | |
| LA6010 | AAGAAAAAGTAATTAATTAATATTATTAAAATTTAATGACAACAGGAGGGCCAC | *PxnanosP* 3'UTR | |
| LA6011 | GTATCTTAACGCGAGTTAATTAATGCAAGGCCAACTGACATCTAGAG | | |
| LA3425 | ATGGACTATAAGGACCACGACGGAG | NLS-aCas9-NLS | |
| LA3426 | TTAATTAATTACTTTTTCTTTTTTGCCTGGCCGGCC | | |
| LA4709 | GACCCGTAAGATCCACCGGATCTAGCTAGAATGAATCGTTTTTAAAATAACAAATCAATTGTTTTATAATATTCG | P10 3'UTR | |
| LA4475 | CCGGCCGTTAACTCGAATCG | | |
| LA4710 | TGAGACCAGGTGGATCACCATGCAGCCGGCCGTTAACTCGAATC | | |
| LA3540 | ATCCCAATGTTACTAACCCCTG | Flanking sequence of *Pxyellow* target site | T7E1 assay and amplification of knock-in cassettes |
| LA3434 | AGCCAACGTCATCCTTGTCAAC | | |
| LA5000 | TTGTATCTAACGTCCTTCGCCTCC | Flanking sequence of *Pxkmo* target site | |
| LA4974 | ATTCCGCCCTACGGAGTAGATG | | |
| LA3438 | gaaattaatacgactcactataggAATTCACCAACAAGAGGGTCgttttagagctagaaa | *yellow*-sgRNA4 template | *In vitro* transcription of sgRNAs |
| LA5002 | gaaattaatacgactcactataggATGAGATACCGTACGATGCGgttttagagctagaaa | *kmo*-sgRNA2 tempalte | |
| LA137 | AAAAGCACCGACTCGGTGCCACTTTTTCAAGTTGATAACGGACTAGCCTTATTTTAACTTGCTATTTCTAGCTCTAAAAC | Common reverse primer | |

| | | | |
|---|---|---|---|
| LA3862 | TGTCGGCATTGAGAGATGGA | 5’ junction of AGG1619 integrated locus | Validation of HDR integration in *Pxyellow* and *Pxkmo* target sites |
| LA2196 | CCAGTTCGGTTATGAGCCGT | | |
| LA1975 | GCGAGTGTATAGCGAGCTAGTG | 3’ junction of AGG1619 integrated locus | |
| LA4998 | TCATGATTCATATACCTCGTAGTTTCCTCTCAAC | | |
| LA4996 | CACCACAAGTATACTGCAGGCAG | 5’ junction of AGG1962 integrated locus | |
| LA4972 | ACCGCAGATAGAAGCGAGAATAG | | |
| LA4973 | AACCTGGCGTTATCTGTGAGGG | Knock-in cassette of AGG1963 | |
| LA323 | ACCAAATCTGCCAGCGTCAATAG | 3’ junction of AGG1962 and AGG1963 integrated locus | |
| LA4975 | TGGCCATCTTCAGTCGGTGG | | |

**Table S2 Germline-specific candidate genes**

| Reference gene | Protein ID | Origin | Homolog in DBM |
|---|---|---|---|
| *Probable ATP-dependent RNA helicase YTHDC2*（*BGCN*） | XP_021711078.1 | *Aedes aegypti* | *Pxbgcn*（g7986.t1） |
| *Inactive peptidyl-prolyl cis-trans isomerase shutdown*（*Shutdown*） | XP_001661808.2 | | *Pxshu1*（g8286.t1）<br>*Pxshu2*（g10629.t1） |
| *Sin3 histone deacetylase corepressor complex component*（*SDS3*） | XP_021712942.1 | | *Pxsds3*（g19605.t1） |
| *Meiotic W68*（*Meiw68*） | AAF57553.1 | *Drosophila melanogaster* | *Pxmeiw68*（g25679.t1） |
| *Silkworm PIWI*（*SIWI*） | BAF98574.1 | *Bombyx mori* | *Pxpiwi*（g34084.t1） |
| *NanosO* | BAF63425.1 | | *PxnanosO*（g1550.t1） |
| *NanosN* | ABS17681.1 | | *PxnanosN*（g16294.t1） |
| *NanosP* | BAF63424.1 | | *PxnanosP*（g60.t1） |
| *NanosM* | BAF73619.1 | | *PxnanosM*（g19168.t1） |

**Table S3 Flanking sequences of *Pxvasa-Cas9* and *Pxmeiw68-Cas9* transgenic lines**

| **Transgenic line** | **5' flanking sequence*** | **3' flanking sequence*** | **Insertion locus** |
|---|---|---|---|
| 1536A | TTTGAATTGGTAGGTAATCGTTTTTACATTAA | TTAACACATACCTATATAGTAGCCCTCTTGCT | Intergenic region |
| 1536C | ATTCTAGTTTCTAAAACAAAAATCGGATTTAA | TTAATAAGCGATAAGAATTGAGGCCCGCTAC | 3'UTR of 3-hydroxy-3-methylglutaryl-coenzyme A reductase (LOC105398578) |
| 1536D | CGGAGTTACCACCACCGAAAATTAGCCATTAA | TTAAGCATCGTCGCTCGCTTATCAAAATCGG | Intron of uncharacterized LOC105380296 |
| 1536E | ATGTGCTTGGCAATGCAATTATAAGTACCTTAA | TTAAAATACTGGTGCAACGTGGATAAAAAA | Intron of histone-lysine N-methyltransferase，H3 lysine-79 specific-like (LOC105387142) |
| 1906A | TATTTACTTACGAAGTTTAATATAAGTACTTAA | TTAACATGGATTAACAAGGTTTAACAGAAAG | Intergenic region |
| 1906B | TGACTAAAATCTGTATGCTGAATTCTATATTAA | TTAACTGTAAACATACAATCATACCAAGTAAT | Intergenic region |
| 1906C | TCTGGGAAGATGAAGAATTGGTAGCTAATTAA | TTAAATGAAGAAATTACGTCAGTCTTCAACC | Intergenic region |
| 1906D | TTTAGGTCGAGTAAAGCGCAAATCTTTTTTAA§ | TTAAATAATAGTTTCTAATTTTTTTATTATTCA§ | Unknown |
| 1906E | TAGGTAAATTCAGCACACATAATATGATTTTAA | TTAAACACATGCTAGTGCTTGTTATCCCTGAA | Unannotated genomic region |
| 1906F | AAGGACAACAAAAGAAGACAGACGATGTTAA | TTAATAATACGTACATAGTAATCGACGTGCTC | Intron of LOC105394090 |
| 1906G | TTTAGGTCGAGTAAAGCGCAAATCTTTTTTAA§ | TTAAATAATAGTTTCTAATTTTTTTATTATTCA§ | Unknown |
| 2093A | ACGTTTTTGCTCATGTTACTGCAGCATCTTTAA | TTAAAAATAAAGTGGTTAATAACTTGGTGAA | Intergenic region |
| 2093B | AAACACGGACCAACGAATCGATTTTTTTTTAA | TTAAATATCCTACAAATGGGATTTGAAATATG | Intergenic region |

| | | | |
|---|---|---|---|
| 2093C | ACTTAACTTACTAGGCTTTTACTCCGTATTTAA | TTAAGGGAACCAACTTTGTACTCGTATAGTTT | Intron of dual 3',5'-cyclic-AMP and -GMP phosphodiesterase 11-like (LOC105390748) |
| 2093D | CGTGGTGCGGAGGGCGGTGGATCTACTTAA | TTAAGAAAGGTATGGCTTCGGGCATTTCATAT | 3’UTR of acylamino-acid-releasing enzyme-like (LOC105388523) |
| 2093E | CAACAGTAACATCGCACGTTTGAATCTTTTAA | TTAAAGCAGTTTTAAGTGGAGTACGGTCGGC | Intron of metalloproteinase inhibitor 3 (LOC105388015) |

*** The TATA insertion site characteristic of *piggyBac*-mediated insertion is underlined. This sequence is duplicated on insertion and therefore appears on both sides of the inserted transgene.**

**§ Backbone sequences of donor plasmid.**
